# Extracellular vesicle microRNA cargo engineering reveals critical mechanisms underlying therapeutic efficacy

**DOI:** 10.1101/2022.01.31.478505

**Authors:** Lindsey M. Euscher, Kyle I. Mentkowski, Touba Tarvirdizadeh, Isabella Julian, Karan Bhatt, Lisa Eagler, Jennifer K. Lang

## Abstract

**Background:** Extracellular vesicles (EVs) are key mediators of intercellular communication and function to transfer biological cargo, including microRNA (miR), from donor to recipient cells. EVs isolated from cardiosphere-derived cells (CDCs) have demonstrated therapeutic efficacy in pre-clinical models of ischemic heart disease, highlighting them as promising vectors for the treatment of CVD. Importantly, it has not yet been established whether miR cargo is necessary for the observed therapeutic benefit of CDC-EVs following acute MI (AMI).

**Methods:** CDCs were transfected with siRNA against Drosha, the initial endonuclease in the miRNA biogenesis pathway, to generate miR depleted DROSHA-EVs. EVs were characterized by size, morphology, and protein/miR expression. The role of EV miRNA on cardiac target cell apoptosis, proliferation and angiogenesis was examined using a series of *in vitro* assays. Mice with acute MI underwent delivery of human CDC EVs, DROSHA-EVs and placebo in a double-blind study. LVEF was assessed by echo at 1- and 28-days post-MI and tissue samples processed for assessment of histological endpoints. *In vitro* sufficiency assays were performed using a combinatorial approach with individual candidate miRs to identify clusters exhibiting synergistic efficacy.

**Results:** DROSHA-EVs exhibited global downregulation of miRNA cargo but were otherwise indistinguishable from wild-type CDC-EVs. miR cargo was responsible for mediating the beneficial effects of human CDC-EV treatment on cardiomyocyte apoptosis, fibroblast proliferation and angiogenesis *in vitro*. DROSHA-EVs were unable to promote recovery following AMI on a functional or histological level, highlighting the critical role of EV miRNAs in cardioprotection following ischemic injury. A potentially therapeutic miR cluster, miR-146a-370-126a, was identified which acted synergistically to reduce cardiomyocyte apoptosis and was sufficient to render inert EVs into therapeutic vectors.

**Conclusions:** These results demonstrate for the first time that miRNAs are required for the regenerative potential of CDC-EVs following AMI and identify a novel miR cluster with therapeutic implications.

## Introduction

Extracellular vesicles (EVs), of which exosomes comprise a subset, have been widely identified as key paracrine mediators of intercellular signaling and effectors of stem-cell mediated cardiac repair (1–3). EVs isolated from specific cell populations, including cardiosphere-derived cells (CDCs), have demonstrated therapeutic efficacy in pre-clinical models of ischemic heart disease, highlighting them as promising vectors for the treatment of cardiovascular disease (4–10). Their mechanism of action has been attributed to the transfer of information between cells. Following cellular uptake, EVs deliver their protein, lipid and RNA content to recipient cells, modulating their fate and behavior (11). While prior studies have demonstrated the effects of individual EV encapsulated signaling molecules on both immune and cardiac cells, there is a lack of knowledge on the bioactivity and necessity of specific classes of endogenous cargo in mediating the efficacy of therapeutic EVs following ischemic injury to the heart.

We hypothesized that extracellular vesicle miRNA cargo is necessary for the therapeutic benefits of CDC-EVs. Accordingly, we have taken a genetic approach to deplete EV miRNA (miR) by shRNA knockdown of Drosha, a key endonuclease in miRNA biogenesis, to elucidate the relative therapeutic contribution of miRNA as a cargo class following myocardial infarction (MI) (12)(13). In this report, we demonstrate that human CDC-EVs depleted of miRNA cargo are unable to recapitulate the cardioprotective effects of wild-type hCDC-EVs, including modulation of cardiomyocyte apoptosis, fibroblast proliferation and angiogenesis in a series of comprehensive *in vitro* assays. *In vivo* studies in a murine model of acute MI (AMI) demonstrate the necessity of CDC-EV miRNA in modulating cardiomyocyte apoptosis, interstitial fibrosis, macrophage density and enhancing systolic function following AMI. Interestingly, we show species-specific differences in therapeutic efficacy between mouse and human CDC-EVs that further underlie the mechanism of action of CDC-EVs. Finally, using a series of sufficiency assays, we demonstrate the synergistic activity of three key miRNAs in mediating the cardioprotective effects of CDC-EVs on cardiomyocyte apoptosis and fibroblast proliferation. These results provide important insight into the mechanism of action of stem cell derived extracellular vesicles in cardioprotection post-MI and highlight a role for miR cargo engineering in translating EV therapeutics from bench to bedside.

## Methods

All animal experiments were performed according to protocols approved by the University at Buffalo Institutional Animal Care and Use Committee (IACUC). Following written consent, human cardiosphere-derived cells were obtained from patients who consented to tissue use under protocols approved by the University at Buffalo Research Subjects Institutional Review Board (IRB) as previously described (14). Methods were carried out in accordance with the relevant guidelines and regulations.

### Statistical Analysis

Quantitative endpoints are summarized with means and SEMs. Qualitative data are summarized with percentages. Differences in quantitative endpoints between treatment groups with one independent variable were assessed using unpaired student t-tests (two groups) or one-way ANOVA and Tukey’s post-hoc test (three or more groups). Three-way ANOVA with Dunnett’s multiple comparisons test was used to determine if there was an interaction effect between three independent variables on a continuous dependent variable. Statistical significance cutoff was set to p<0.05.

### EV Isolation and Characterization

EVs were isolated from human and mouse cardiosphere-derived cells (CDCs) using an ultrafiltration-based method as previously described (15, 16). Briefly, supernatants were first collected from either human or mouse CDC culture flasks, passed through a 0.2μm filter, and subsequently concentrated using an Amicon Ultra-15 centrifugal filter (Millipore Sigma) with a 10kDa molecular weight cutoff. Isolated extracellular vesicles were characterized by size using Nanoparticle Tracking Analysis (NTA), morphology by cryo-TEM (FEI Vitrobot, Mark IV), and expression of known exosomal surface protein markers and markers of cellular contamination (Exo-Check Exosome Antibody Array, SBI) as previously described (15, 16). Protein content was quantified using a Pierce BCA Protein Assay Kit (Thermo Fisher Scientific) following manufacturer’s protocols and used as a surrogate measure of EV concentration for downstream applications.

### Cell Culture

#### Cardiosphere-Derived Cells

Human and mouse CDCs were generated and characterized as previously described following protocols approved by the University at Buffalo IRB and IACUC (14, 17). Briefly, human endomyocardial biopsies from the right atrial appendage or murine atrial tissue from adult C57BL/6 wild type mice were minced into small fragments, washed, and subsequently digested using type IV collagenase for 60 minutes at 37°C. Explants were then cultured on 20μg/mL fibronectin-coated tissue culture plates. A layer of stromal-like cells, and small, round, phase-bright cells migrated outwards and surrounded the explant. Once confluent, cells surrounding the explants were harvested using gentle enzymatic digestion. These cardiosphere-forming cells were seeded on low attachment dishes and cultured in Iscove’s modified Dulbecco medium containing 20% heat inactivated fetal bovine serum, penicillin/streptomycin 100μg/ml, 2mmol/L L-glutamine and 0.1mmol/L 2-mercaptoethanol. Following 4-6 days in suspension culture, free-floating cardiospheres were harvested by aspiration and seeded onto fibronectin coated tissue culture flasks, forming monolayer cardiosphere-derived cells. Cells were characterized by flow cytometry and immunohistochemistry with hematopoietic, mesenchymal and cardiac markers as described previously (14).

#### Bone Marrow Derived Macrophages

Femurs were isolated from M/F 3- to 4-month-old C57BL/6 mice. The cellular fraction of the bone marrow was isolated by flushing PBS through the medullary cavity and subsequent filtration of the isolate through a 70μm mesh filter. Cells were resuspended in RPMI 1640 media supplemented with 10% FBS, 1% penicillin/streptomycin and 30ng/mL of M-CSF (Sigma Aldrich) and subsequently plated at a density of 4×10^5^ cells/mL. After two days in culture, additional media was added to the wells, with a complete media change performed on day 4 and repeated every other day thereafter. Starting on day 5, this population was referred to as naïve BMDMs. On day 6 of BMDM culture, cells were activated for 24 hours by 100ng/mL LPS (Sigma Aldrich) and 50ng/mL IFNγ (R&D Systems) towards an M1 like phenotype. 24 hours after polarization, cells were treated with 10μg of EVs per 2×10^5^ cells for 24 hours prior to PCR analysis.

#### Primary Neonatal Cardiomyocytes

Primary neonatal cardiomyocytes were isolated from C57BL/6 mouse neonates on postnatal days 3-5 using the Pierce Primary Cardiomyocyte Isolation Kit (Thermo Fisher) as previously described (14). Media was changed 24 hours after isolation and every other day thereafter. On day 6 of culture, cells were treated with 10μg of EVs per 4×10^5^ cells for 24 hours prior to fixation and permeabilization.

#### Normal Human Dermal Fibroblasts

Normal Human Dermal Fibroblasts (NHDFs) (ATCC) were cultured in RPMI 1640 Media, 10% FBS, 1% penicillin/streptomycin. Cells were treated with 10μg of EVs per 2×10^5^ cells for 24 hours prior to fixation and permeabilization.

#### HUVECs

HUVECs (Thermo Fisher) were cultured on rat tail collagen I coated plates in EBM2 Endothelial Growth Medium (Lonza). Cells were treated with 10μg of EVs per 3×10^5^ cells for 24 hours prior to angiogenesis and migration assays. All cells were incubated at 37°C in 5% CO_2_.

### Drosha siRNA Knockdown

Human or mouse CDCs were transfected with small interfering RNAs against Drosha (Thermo Fisher HSS178992 or MSS274198) or a Medium GC Content Negative Control (Thermo Fisher) using Lipofectamine RNAiMAX (Thermo Fisher) or TransIT TKO (Mirus) following manufacturer’s protocols. BLOCK-iT Fluorescent Oligo (Thermo Fisher) was used as a loading control. Following a 24-hour incubation, media was changed to remove the transfection complex. For mouse CDCs, media was changed again on day 3 post transfection to serum free media and mDrosha-EVs isolated on day 10 post transfection. For human CDCs, media was changed on day 5 to serum free media and hDROSHA-EVs isolated on day 12 post transfection.

### miR Mimic Transfection

Murine neonatal cardiomyocytes and NHDFs were transfected with miR-146a-5p, miR-126-3p and miR-370-3p mimics (Lightswitch, 100nM final concentration) using Trans-IT TKO (Mirus) in their respective serum free growth media. To generate miR loaded NHDF-EVs, the media was changed at 24 hours post-transfection to remove the transfection complex and the cells left undisturbed for an additional 7 days. After a week, the supernatant was collected and EVs isolated and characterized as described above.

### Quantitative Real-Time PCR

RNA was extracted from cells using an E.Z.N.A. Total RNA Kit I (Omega Bio-tek). cDNA was synthesized using SuperScript III reverse transcriptase (Thermo Fisher). Quantitative real time PCR was performed using SsoFast EvaGreen Supermix (Bio-Rad) or PowerUp SYBR Green Master Mix (Thermo Fisher) for Drosha (**Table I**). Samples were run in triplicate and gene expression calculated by ΔΔC_t_ analysis using 18S as a reference. Statistical significance was tested on log_2_-transformed data using unpaired student’s t test.

**Table I.**
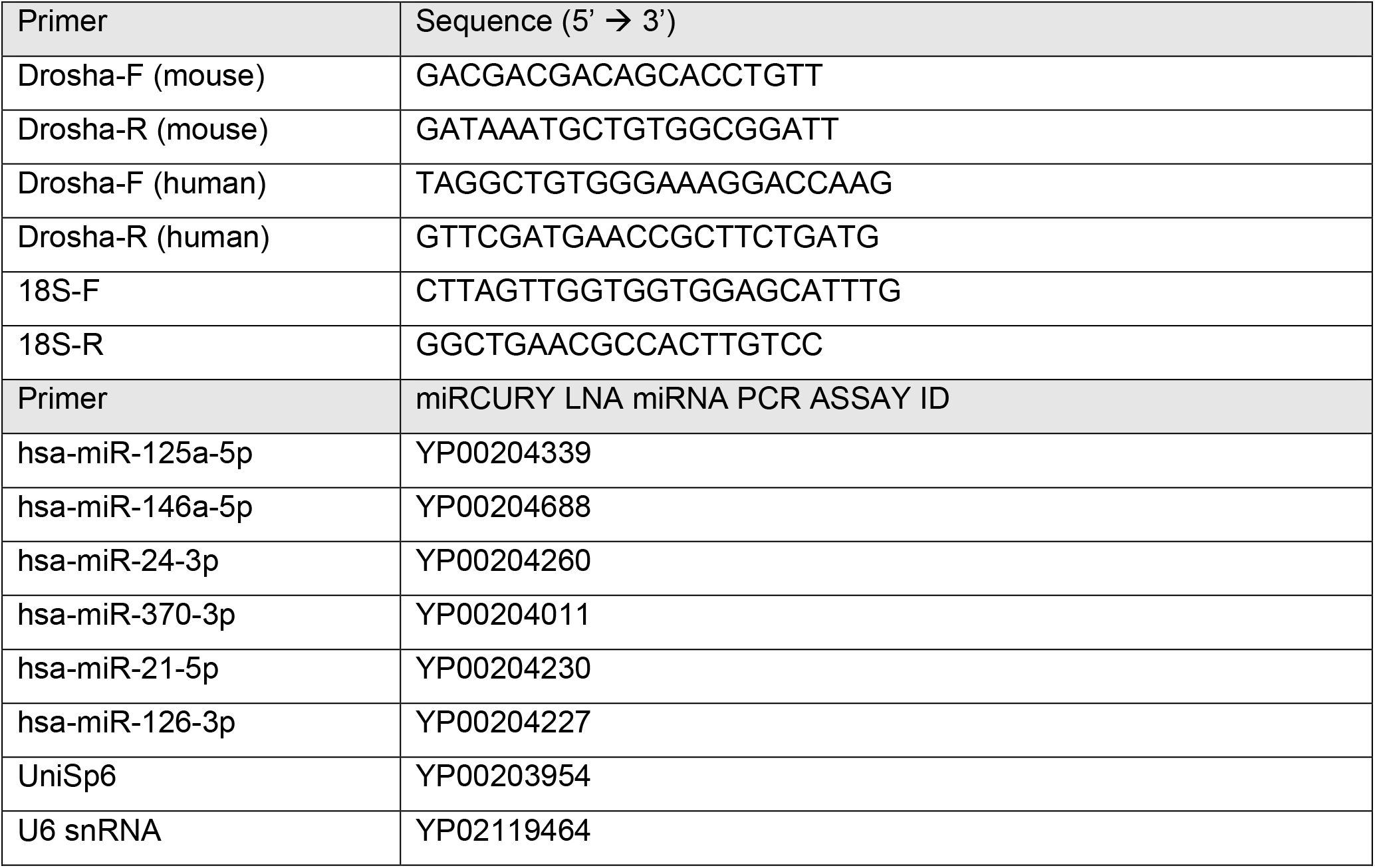
Primers used for RT-PCR.

### miR RT-PCR

EV samples were diluted 1:1 in QIAzol lysis reagent and sonicated using a constant duty cycle, power 3 and 3 pulses per sample, resting on ice between pulses. Samples were then allowed to incubate at room temperature for 5 minutes. Subsequent RNA isolation was performed using miRNeasy Mini Kit (Qiagen) and following the manufacturer’s protocol (Purification of Total RNA from Animal Cells, miRNeasy Mini Handbook). RT-PCR for specific miRNAs was performed using miRCURY LNA miRNA PCR Kit (Qiagen) following the manufacturer’s protocol (miRCURY LNA miRNA PCR-Exosomes, Serum/Plasma and Other Biofluid Samples Handbook) (**Table I**). For PCR performed on cell lysates, RNA was extracted from cells using an E.Z.N.A Total RNA Kit I (Omega Bio-tek) and subsequent cDNA and PCR was run using the miRCURY LNA miRNA PCR Kit (Qiagen) following manufacturer’s protocols.

### Cardiomyocyte Apoptosis Assay

Primary cardiomyocytes were treated with 10μg of EVs per 4×10^5^ cells for 24 hours prior to fixation with 4% PFA for 15 minutes and permeabilization for 20 minutes at room temperature. Subsequent TUNEL assay for apoptosis was performed using Click-iT TUNEL Alexa Fluor 488 Imaging Assay (Thermo Fisher) following manufacturer’s protocol. Cells were then incubated with Cardiac TnI primary antibody (Rabbit polyclonal, Proteintech) at 1:100 dilution overnight at 4°C. Donkey anti-rabbit secondary antibody (Thermo Fisher) was incubated with cells at a 1:400 dilution for 1 hour at room temperature in the dark. DAPI (1X) was incubated with cells for 5 minutes for nuclear staining followed by PBS washing and imaging. Images were obtained using an EVOS FL Auto Imaging System. Apoptosis was assessed as the percent of TUNEL positive cardiomyocytes.

### Fibroblast Proliferation Assay

NHDFs were treated with 10μg of EVs per 2×10^5^ cells for 24 hours prior to fixation with 4% PFA for 15 minutes and permeabilization for 20 minutes at room temperature. A subsequent EdU Assay for proliferation was performed using Click-iT EdU Cell Proliferation Kit for Imaging (Thermo Fisher) following the manufacturer’s protocol. DAPI (1X) was incubated with cells for 5 minutes for nuclear staining followed by PBS washing and imaging. Images were obtained using an EVOS FL Auto Imaging System. Proliferation was assessed as the percent of EdU positive fibroblasts.

### HUVEC Angiogenesis Assay

HUVECs were treated with 10μg of EVs per 3×10^5^ cells for 24 hours prior to seeding. 50μL of ice cold Matrigel was loaded into a pre-chilled 96 well plate using chilled pipette tips and allowed to polymerize at 37°C for 1 hour. HUVECs were seeded at 5,000 cells/well and imaged at 10X after a 4-hour incubation using an EVOS FL Auto fluorescence microscope.

### HUVEC Migration Assay

HUVECs were seeded at 40,000 cells/well on a 48 well plate pre-coated with rat tail collagen approximately 18 hours prior to scratching. Immediately after the scratch, cells were treated with 10μg of EVs per 3×10^5^ cells. Wells were imaged at 0, 2, 4 and 6 hours after scratch using an EVOS FL Auto fluorescence microscope at 10X magnification. Fiji was used to calculate the change in scratch area at each time point compared with the initial scratch.

### Murine Model of Acute MI

#### Thoracotomy and LAD ligation

Anesthesia was induced in adult 8-12wk C57BL/6J mice with a mixture of ketamine (100 mg/mL) and xylazine (20 mg/mL). A midline ventral cervical skin incision was performed allowing the muscles overlying the larynx and trachea to be bluntly dissected and retracted for visualization of the intubation device through the exposed trachea. The tongue was slightly retracted and a 20-gauge 1” smooth needle was inserted through the mouth and larynx and into the trachea for intubation. The needle was advanced 8 to 10 mm from the larynx and taped in place to prevent dislodgement. Sterile lubricating drops were placed in the eyes. Mice were ventilated with room air supplemented with oxygen (1 L/min) at a stroke rate of 130 strokes/min and a tidal volume of 10.3 μL/g using an Inspira ASV small animal ventilator (Harvard Apparatus, Germany). The left chest was shaved, and a 1.5 cm skin incision made along the mid-axillary line. The left pectoralis major muscle was bluntly dissected and retracted, and a left thoracotomy performed between the third and fourth ribs to visualize the anterior surface of the heart and left lung. The LAD was visualized emerging from under the left atrium and ligated with an 8-0 nylon suture. Successful infarction was verified by visualization of blanching of the distal myocardium and dynamic ECG changes (Rodent Surgical Monitor+, Indus Instruments). Using 6–0 polypropylene suture, the intercostal space was closed in an x-mattress pattern, followed by closure of the muscle layer in an interrupted suture pattern, and skin layers in a running mattress pattern. Mice received post-operative analgesia with 3.25 mg/kg SQ extended-release buprenorphine (Ethiqa XR, Fidelis Pharmaceuticals) and were recovered on a heating pad.

#### Intramyocardial injection

Immediately following LAD ligation, mice were randomly selected to receive intramyocardial injections of PBS, CDC EVs, or DROSHA EVs. Investigators were blinded to treatment groups. Equal numbers of male and female mice were selected across groups to control for sex as a variable. Mice received a total of 100μg of EVs (or matched volume vehicle control) administered intramyocardially in two peri-infarct sites (10μl/injection) using a Hamilton syringe with a 30.5-gauge sterile beveled needle. The tip of the needle was bent at a 45° angle and introduced into the left ventricular myocardium. The solution in the syringe was slowly injected and local tissue blanching verified, indicating intramyocardial delivery. The syringe was held in place for an additional 3–5 seconds before withdrawal.

### Echocardiography

Transthoracic echocardiography (Vivid E9 with GE Ultrasound i13L Intraoperative Epicardial Probe) was performed on mice 2 days and 28 days post-MI. Depilatory cream was applied to the chest to remove hair to reduce imaging artifacts. Mice were placed on a heated surgical platform (Rodent Surgical Monitor+, Indus Instruments) to maintain appropriate body temperature and noninvasively monitor heart rate, continuous ECG, respiration rate, and SpO_2_ during imaging. LVEF was calculated from M-mode images obtained in the parasternal short axis (PSAX) mid-papillary view. Heart rate was maintained between 500 and 550 bpm during image acquisition. Investigators were blinded to treatment groups at the time of image acquisition and data analysis.

### Tissue Processing

Animals were euthanized at the terminal endpoints of the study and hearts perfused with KCL followed by PBS. A portion of hearts were fixed for 30 min in 4% paraformaldehyde (PFA), cryopreserved in sucrose gradient (6, 15%) for 24 hours, snap frozen in OCT (optimal cutting temperature) compound (Tissue-Tek) and sectioned transversely at 12μm thickness using a Leica cryostat. Additional hearts were fixed in 10% normal buffer formalin (NBF) overnight at room temperature, sectioned transversely into 1mm thick slices, and paraffin embedded apex to base. Paraffin blocks were further transversely sectioned at 5μm thickness and stained with Trichrome by the University at Buffalo Histology Core.

### Immunohistochemistry

OCT sections were permeabilized for 15 minutes with 1% Triton X (Thermo Fisher) and blocked for 2 hours with a solution containing 0.5% TritonX-100 and 5% donkey serum (Thermo Fisher). Primary antibodies used were rat CD68 (FA-11, 1:100; Thermo Fisher), rabbit cardiac troponin I (1:50; Proteintech), F-actin (1:200; Rhodamine Phalloidin, Thermo Fisher) and rabbit cleaved caspase-3 (1:200; Asp175, Cell Signaling). Alexa Fluor 488-, 555-, and 647-conjugated secondary antibodies (Thermo Fisher) were used at 1:500. Tissue sections were counterstained with DAPI (Thermo Fisher). Images were captured at 20x magnification using an EVOS FL Auto Imaging System and an Olympus IX83 microscope with wide field fluorescence and at 63x magnification using a Leica TCS SP8 Confocal microscope. Macrophage density was calculated using a Fiji thresholding technique as the CD68+ area/total DAPI+ area. The percentage of apoptotic cells was calculated using Fiji as cleaved caspase-3+ cells/total cells. Paraffin embedded trichrome stained sections were used for quantification of interstitial fibrosis. Interstitial fibrosis was determined using a Fiji thresholding method to compare the amount of blue to red staining. All quantification was performed by an investigator blinded to the identity of the samples.

### Data Availability

All data associated with this study are included in the published manuscript.

## Results

### Human CDCs transfected with Drosha siRNA generate miR-depleted extracellular vesicles

To determine if miRNA cargo is necessary for mediating the therapeutic benefit of human CDC-derived EVs, cardiosphere-derived cells were transfected with siRNA against DROSHA, the initial endonuclease in the miR biogenesis pathway, to generate miR-depleted hCDC-EVs (DROSHA EVs) (**Fig 1A**). Transfection of CDCs with siRNA against DROSHA reduced DROSHA mRNA expression in CDCs by 90% (**Fig 1B**), and resulted in reductions of miRNA cargo (as assessed by surrogate miRs) in both CDCs and their EVs by 81±5% and 65±4%, respectively, as compared with Scr control (mean ± SEM, *p<0.001 using unpaired student’s t-test) (**Fig 1C,D**). DROSHA siRNA transfected CDCs demonstrated no change in viability when compared with Scrambled control (Scr) siRNA CDCs at the time of EV collection as assessed by calcein AM staining (**Fig 1E**). miR cargo depletion did not affect the physical properties of CDC-derived EVs (unmodified CDC-EVs previously characterized in (14, 15)) as nanoparticle tracking analysis of Drosha EVs showed a typical size distribution (mean diameters, 78 nm ± 20 nm) (**Fig 1F**) and cryo-TEM demonstrated the expected exosomal morphology of small, round vesicles with a clearly discernible lipid bilayer (**Fig 1H**). DROSHA EVs were positive for known exosomal markers CD63, CD81, ALIX, FLOT1, ICAM1, EpCam, ANXA5 and TSG101 with preparations negative for any cellular contamination (by cis-Golgi marker GM130) (**Fig 1G**). The total amounts of protein cargo were not altered by ablation of miRNA cargo (BCA assay on matched samples, n=8, unpaired student’s t-test, p>0.05). Taken together, DROSHA KD was capable of largely depleting the miRNA content of CDC EVs while not affecting other aspects of EV structural function.

**Figure 1.**
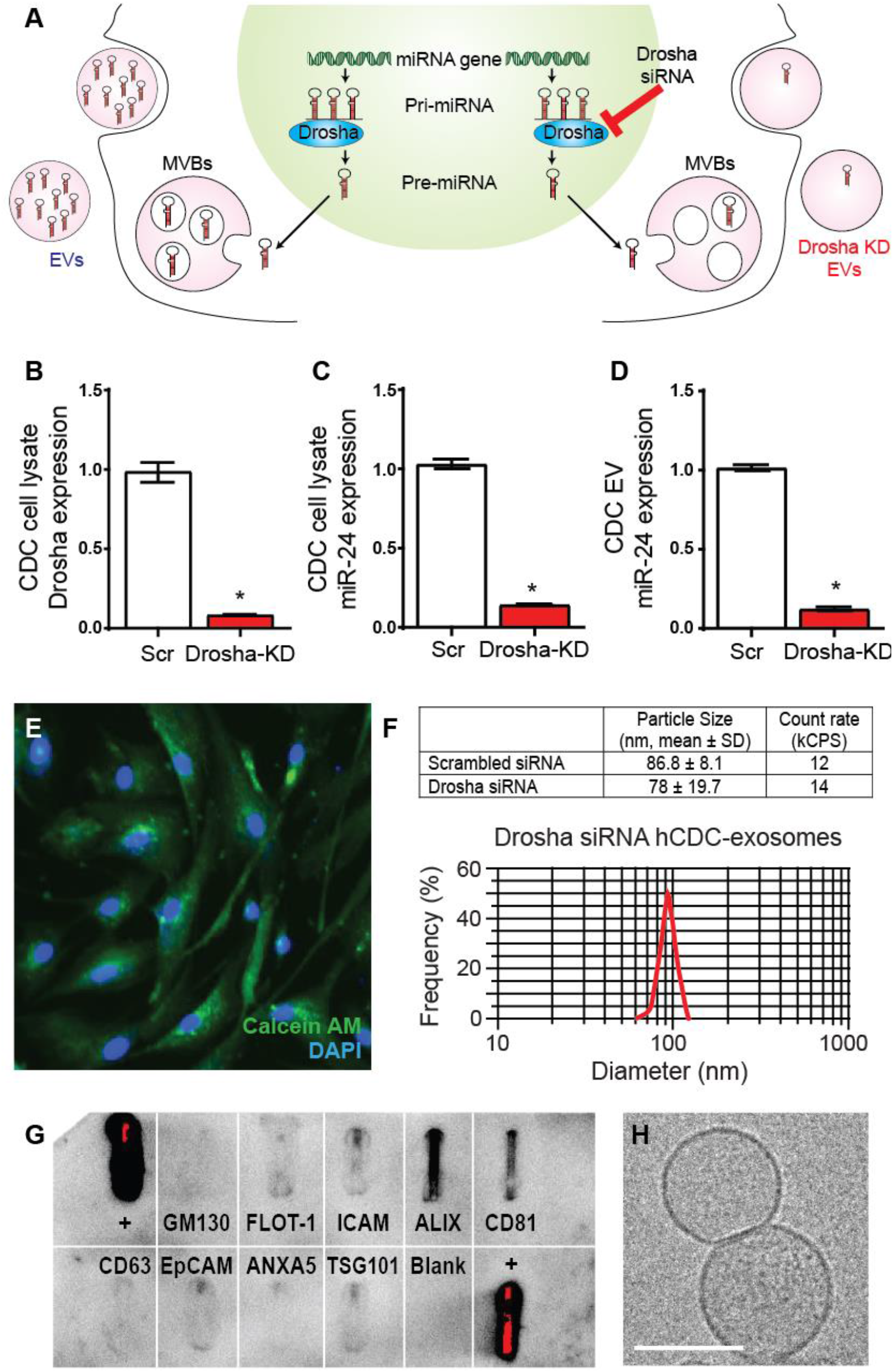
Characterization of DROSHA siRNA transfected human CDCs and their miR depleted extracellular vesicles. **(A)** Schematic of miR biogenesis and miRNA incorporation into EVs. **(B)** Relative expression of DROSHA mRNA in hCDCs transfected with Scr or Drosha siRNA. Relative expression of candidate miRNA (miR-24) in **(C)** hCDCs transfected with Scr and DROSHA siRNA and **(D)** CDC EVs isolated from Scr and DROSHA siRNA transfected cells (n=3). *p<0.0001 using unpaired student’s t-test. Data represented as mean ± SEM. **(E)** Representative image of calcein green AM-stained DROSHA siRNA transfected CDCs at the time of EV collection. **(F)** Scrambled (Scr) and DROSHA siRNA (DROSHA KD) CDC EV size distribution by nanoparticle tracking analysis with representative histogram plot. **(G)** Exo-Check Exosome Antibody Array for exosomal surface protein marker detection and assessment of cellular contamination of DROSHA KD EVs. **(H)** Cryo-TEM image of DROSHA KD extracellular vesicles isolated from serum free tissue culture. Scale bar: 50μM

### miRNA cargo mediates hCDC-EV cardioprotective functions *in vitro*

#### Cardiomyocyte apoptosis

We have previously demonstrated that EVs released from human cardiosphere-derived cells are responsible for inhibiting cardiomyocyte apoptosis *in vitro* (14). To determine if EV miRNA cargo is necessary in mediating human CDC (hCDC) EV suppression of cardiomyocyte apoptosis, we incubated primary mouse neonatal cardiomyocytes with hCDC EVs (EVs) and DROSHA EVs. hCDC EVs, with normal levels of miRNA cargo, had a significant effect on preventing myocyte cell death from apoptosis as assessed by the percentage of TUNEL-positive nuclei at one week when compared with PBS control (23 ± 2% vs. 35 ± 3%, respectively) (**Fig 2A,C**). DROSHA EVs were unable to recapitulate these effects (33 ± 4%) (**Fig 2A,D**), suggesting CDC miRNA cargo was responsible for mediating the protective effects of CDC EVs on cardiomyocyte cell death *in vitro* (n=8, mean ± SEM, *p < 0.05 using one-way ANOVA with Tukey’s post-hoc test).

**Figure 2.**
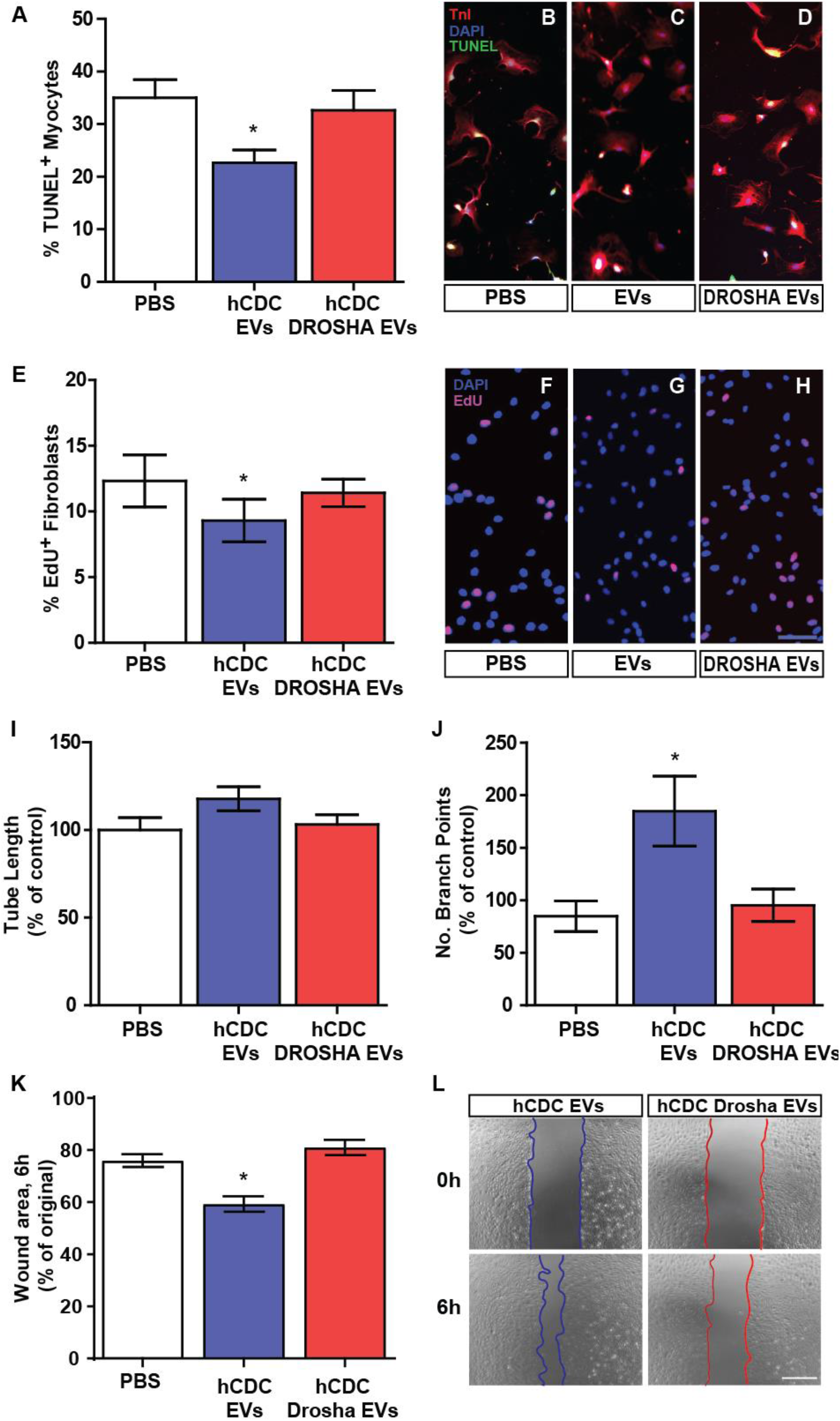
hCDC EV miR cargo mediates cardiomyocyte apoptosis, fibroblast proliferation, and angiogenesis *in vitro*. DROSHA KD CDC EVs were unable to recapitulate control CDC EVs in their ability to **(A-D)** reduce terminal deoxynucleotidyl transferase nick end labeling (TUNEL)-positive cardiomyocytes (CMs) in an *in vitro* apoptosis assay (n=8), **(E-H)** reduce fibroblast proliferation as assessed by EdU staining with a 5-hour pulse (n=4), and **(I-L)** stimulate endothelial migration and angiogenesis *in vitro* (n=7), when compared with cells treated with PBS control (n=7). *p<0.05 using one-way ANOVA with Tukey’s post-hoc test. Data represented as mean ± SEM.

#### Fibroblast proliferation

To determine if EV miRNA cargo is necessary for hCDC EV effects on fibroblast proliferation *in vitro* (14), normal human dermal fibroblasts (NHDFs) were treated with PBS control, hCDC EVs or DROSHA EVs for 24 hours. While hCDC EV treatment of NHDFs resulted in a small but significant reduction in proliferation as quantified by EdU uptake following a 5-hour pulse (9 ± 1%), DROSHA EVs showed no appreciable change when compared with PBS control (PBS: 12 ± 2%, DROHSA-EVs: 11 ± 1%), suggesting hCDC EV miRs were necessary in mediating the anti-proliferative effects of CDC EVs on fibroblasts *in vitro* (**Fig 2E-H**). (n=4, mean ± SEM, *p < 0.05 using one-way ANOVA with Tukey’s post-hoc test).

#### Angiogenesis and endothelial cell migration

Our prior studies have shown that hCDC EVs are capable of inducing angiogenesis and endothelial migration of HUVECs *in vitro* (14). To assess if EV miRNA cargo modulates CDC EV functional effects on endothelial cells, we assessed *in vitro* endothelial tube formation and migration of HUVECs following treatment with CDC EVs or DROSHA EVs. HUVECs treated with CDC EVs for 24 hours showed an increased number of branch points (185 ± 33%) as compared with HUVECS treated with DROSHA EVs (95 ± 15%) and PBS control (n=7, mean ± SEM, *p < 0.05 using one-way ANOVA with Tukey’s post-hoc test) (**Fig 2I**) and a trend towards increased tube length in HUVECs treated with CDC EVs (118 ± 7%) as compared with DROSHA EVs (103 ± 5%) (p=0.14 using one-way ANOVA with Tukey’s post-hoc test) (**Fig 2J**), indicative of enhanced angiogenesis in HUVECs exposed to CDC EV microRNA. In addition, HUVECs treated with CDC EVs demonstrated a smaller wound area at 6 hours in a 2D *in vitro* wound-healing (scratch) assay when compared with HUVECs treated with DROSHA EVs and PBS (CDC EVs: 59 ± 3%, DROSHA EVs: 81 ± 3%, PBS: 76 ± 2%) (**Fig 2K,L**), highlighting the role of CDC EV miRs in endothelial cell migration (n=6-10, mean ± SEM, *p < 0.05 using one-way ANOVA with Tukey’s post-hoc test). Thus, our cumulative results show that miR cargo were responsible for mediating the beneficial effects of CDC-EV treatment previously observed *in vitro*.

### Mouse CDC EVs fail to recapitulate the *in vitro* cardioprotective effects of human CDC EVs likely due to lack of shared therapeutic miR cargo

EV cargo content is known to be specific to the parent cell source and environmental conditions from which EVs originate (18). To ascertain any species-specific differences in therapeutic efficacy between human and murine CDC EVs, we derived mouse cardiosphere-derived cells (mCDCs) from atrial biopsies of wild type male and female 8-12wk C57BL/6 mice, as previously described (17, 19), and isolated and characterized their EVs using established protocols (14–16). Mouse CDCs (mCDCs) exhibited a similar morphology, rate of proliferation, and mesenchymal surface marker expression profile to their human counterparts (**Sup Fig 1A,B**). Similar to human CDC EVs, mCDC EVs demonstrated a comparable size profile by NTA (95 ± 19.5nm; mean ± SD) (**Sup Fig 1E**), an expected EV morphology by cryo-TEM (**Sup Fig 1C**), and expression of known EV markers with no evidence of cellular contamination (**Sup Fig 1D**). Also consistent with our human studies, transfection of mCDCs with siRNA against Drosha reduced Drosha mRNA expression in mCDCs and resulted in reductions in miRNA quantity (as assessed by surrogate miRs) in both CDCs and their EVs (n=8-10, *p<0.0001 by unpaired student’s t-test) without altering protein cargo, as measured by pierce BCA protein assay on matched samples (n=8, unpaired student’s t-test, p>0.05) (**Sup Fig 2A-C**). In contrast to hCDC EVs, however, mCDC EVs failed to elicit any effect on cardiomyocyte apoptosis, fibroblast proliferation, endothelial cell migration or angiogenesis *in vitro* (n=6, p>0.05 using one-way ANOVA) (**Sup Fig 2D-F**). Concordantly, miR-depleted mCDC Drosha EV treatment resulted in a similar rate of myocyte apoptosis, EdU+ fibroblasts and endothelial tube length as treatment with mCDC EVs and PBS (**Sup Fig 2D-F**). Mouse CDC EV treatment of HUVECs did result in a decrease in branch points in a 4-hour Matrigel *in vitro* tube assay (**Sup Fig 2G**).

Based on these observations, we hypothesized that the factor/s responsible for divergence of *in vitro* therapeutic efficacy between human and mouse CDC EVs was due to species-specific differences in their therapeutic miR cargo. Using miRNA qPCR, we compared expression levels of several highly expressed microRNAs that have been previously implicated in cardioprotective responses. Consistent with our hypothesis, we found that these miRs were highly expressed in human CDC EVs but were either absent or 1000-100,000x fold less prevalent in their murine counterparts (n=3, *p<0.0001 using an unpaired student’s t-test) (**Sup Fig 3**). This further highlights the significance of EV miRNA as a therapeutic cargo class and underlies the importance of understanding species-specific differences amongst matched EV populations.

### miRNA cargo is necessary for hCDC-EV mediated augmentation of cardiac function following acute MI

To determine whether miR cargo were indeed necessary for the therapeutic benefits afforded by hCDC EVs, CDC EVs and DROSHA EVs were administered by intramyocardial injection at the time of MI and followed with serial echocardiography to assess changes in left ventricular ejection fraction (**Fig 3A**). At 1-day post-MI, both EV groups and PBS vehicle control demonstrated equivocal LV systolic function (**Fig 3B**). At 28 days post-MI, while intramyocardial administered CDC-derived EVs improved LVEF compared with vehicle control treated animals (62 ± 4% vs 44 ± 4%), mice treated with DROSHA EVs demonstrated no significant difference in LVEF (42 ± 6%) compared with vehicle control (n=8 mice/group, mean ± SEM, p<0.05, one-way ANOVA followed by Tukey’s post-hoc test) (**Fig 3B**). This was associated with a trend in decreased infarct size in CDC EVs (23 ± 5.2%) compared with PBS-treated controls (35 ± 6.5%) and DROSHA EVs (33 ± 4.7%) when analyzed by endocardial length-based measurement of Masson’s Trichrome stained mid LV cavity sections from PBS control, CDC EV and DROSHA EV treated hearts at 28 days post-MI (n=6-7 mice/group, mean ± SEM, p=0.10 using one-way ANOVA followed by Tukey’s post-hoc test) (**Fig 3C-D**).

**Figure 3.**
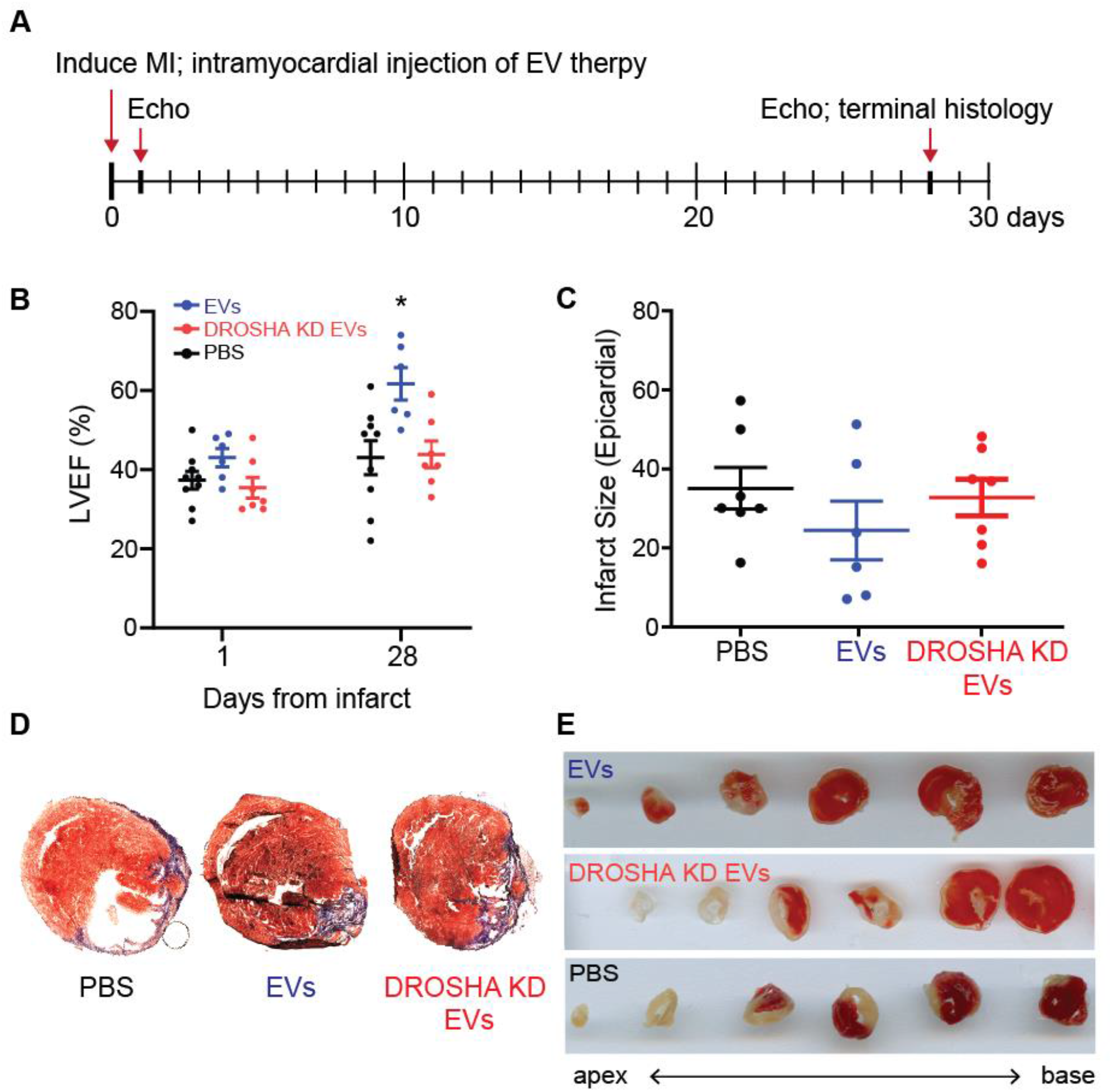
Extracellular vesicle miR cargo is necessary for hCDC EV mediated cardioprotection post MI. **(A)** Schematic depicting the timeline of MI induction and treatment with DROSHA KD EVs and control CDC EVs. **(B)** Left ventricular ejection fraction (%) at 1- and 28-days post MI of mice treated with saline, DROSHA EVs and control CDC EVs demonstrated no difference between DROSHA KD EV and saline groups (n=6-8 mice per group). **(C)** Infarct size quantified by length in all three treatment groups 28 days after injury (n=6-8 mice per group). **(D)** Representative Masson trichrome staining of mid-LV cavity heart sections and **(E)** TTC staining of sequential 0.5mm sections of cardiac tissue at the terminal study endpoint in mice treated with PBS, DROSHA KD EVs and control EVs. (n=6-8 mice per group). *p<0.05 using one-way ANOVA with Tukey’s post-hoc test. Data represented as mean ± SEM.

### hCDC EV miR cargo regulates remote interstitial fibrosis following ischemic injury

Given the difference in therapeutic efficacy between hCDC EVs and miR-depleted DROSHA EVs post-MI, we focused on an immunohistological analysis of infarct and remote myocardium to identify any cellular changes between treatment groups mediated by EV miR cargo. To assess the effect of CDC EV miRs on remote zone interstitial fibrosis, we analyzed the remote regions of Masson’s trichrome stained LV sections at 28 days post-MI. While CDC EVs demonstrated a significant reduction in interstitial fibrosis when compared with PBS control (5 ± 0.3% vs. 10 ± 1%), DROSHA EVs had no effect on reactive fibrosis at 28 days post-MI (8 ± 1%) (n=3-5 mice/ group, mean ± SEM, *p<0.05 using one-way ANOVA followed by Tukey’s post-hoc test) (**Fig 4A-D**). These results highlight the necessity of CDC EV miR cargo in mediating reactive fibrosis following AMI.

**Figure 4.**
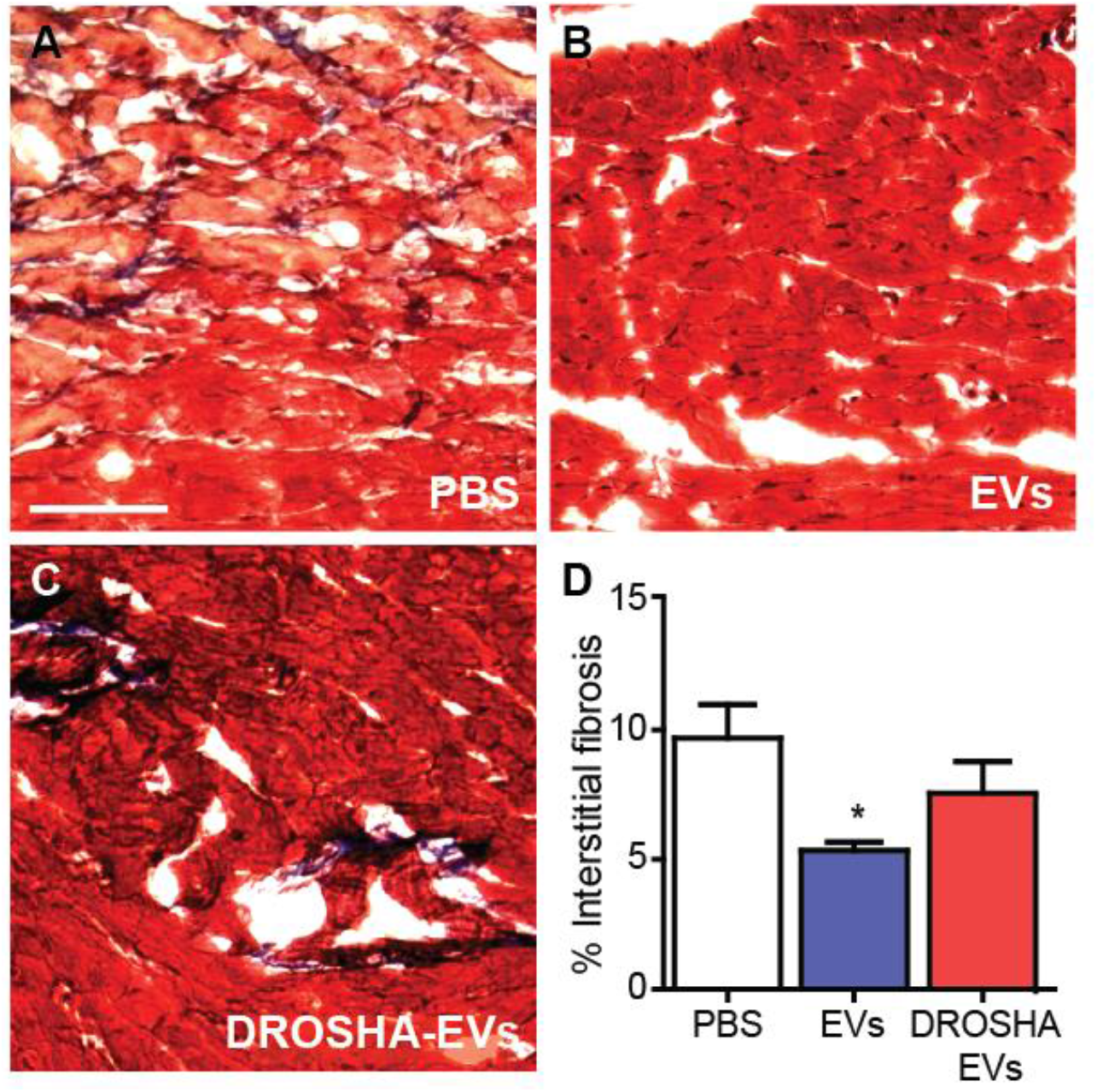
hCDC EV miR cargo regulates reactive interstitial fibrosis follow ischemic injury. Representative trichrome-stained sections from the LV remote region of **(A)** PBS, **(B)** CDC EV and **(C)** DROSHA KD EV treated mice at terminal study endpoint. (**D**) While control CDC EVs demonstrated a reduction in interstitial fibrosis in the remote region 28 days post-MI, DROSHA KD EVs showed no significant difference in reactive fibrosis when compared with PBS vehicle control (n=3 mice per group). Scale bar = 100 μm. *p<0.005 using a one-way ANOVA with Tukey’s post-hoc test. Data represented as mean ± SEM.

### miR cargo is responsible for CDC EV modulation of macrophage density in the ischemic and remote myocardium post-MI

To investigate the effect of EV miRNA on regulating macrophage infiltration following ischemic injury, we analyzed CD68^+^ macrophage density in the infarct and remote zones 28 days post-MI. Macrophage density was highest in the infarct region and lowest in the remote zone for all treatment groups at one month. CDC EV treatment resulted in a significant reduction in CD68+ macrophage density in the infarcted (CDC EV: 6 ± 1%, PBS: 9 ± 1%, DROSHA EV: 9 ± 1%) (**Fig 5A-C,G**) and remote myocardium (CDC EV: 1 ± 0.2%, PBS: 2 ± 0.2%, DROSHA EV: 2 ± 0.1%) (**Fig 5D-F,H**) when compared with PBS control. This effect was dependent on miR cargo as DROSHA-EVs treatment did not have a significant effect (n=3-4 mice/group, mean ± SEM, *p<0.05 using one-way ANOVA followed by Tukey’s post-hoc test).

**Figure 5.**
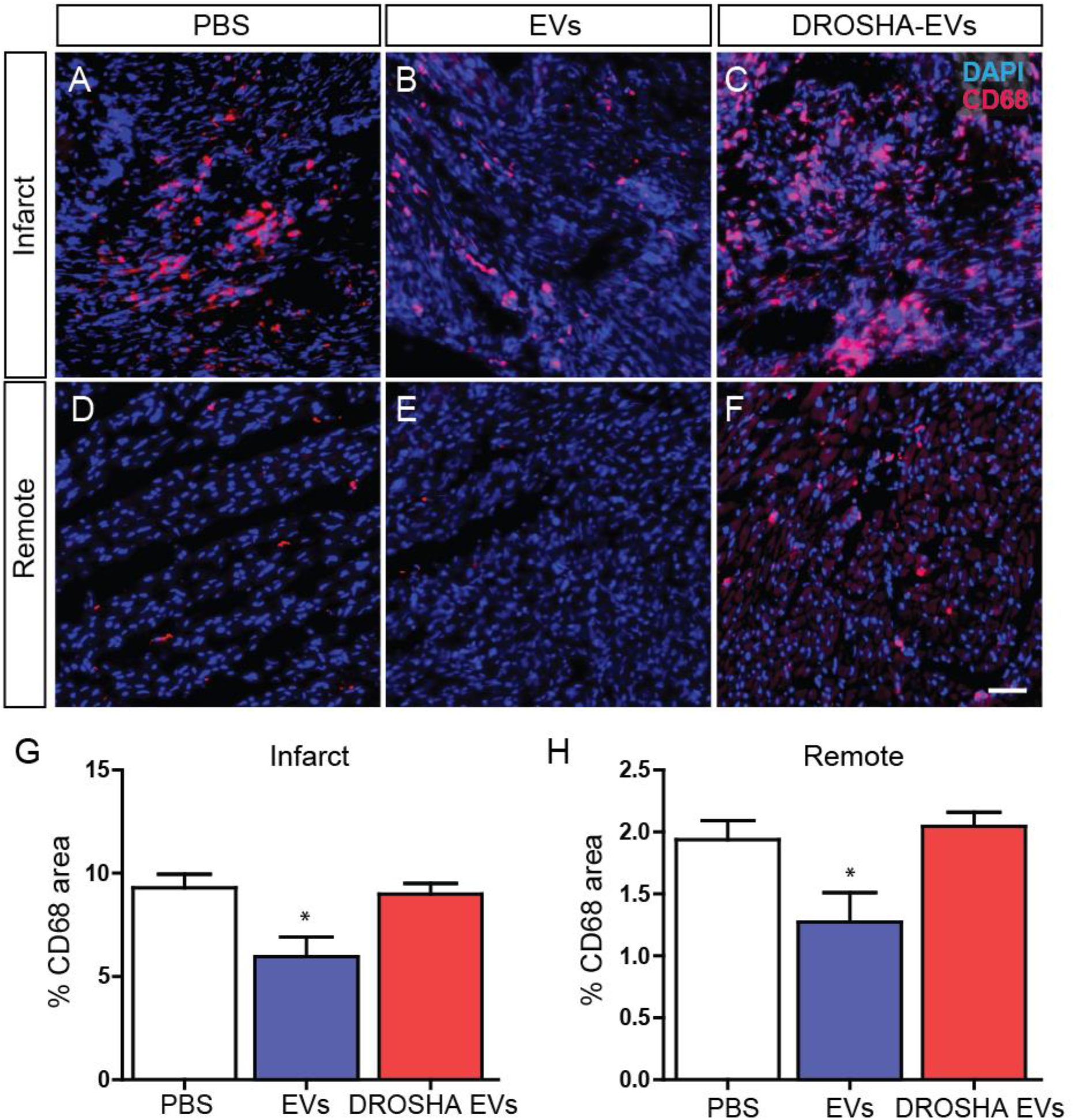
hCDC EV miR cargo regulates macrophage density in ischemic and remote myocardium. CD68-stained sections from the **(A-C)** infarct and **(D-F)** remote zone 28 days post-MI. DROSHA KD EVs failed to recapitulate the reduction in CD68+ macrophage density in the **(G)** infarct region and **(H)** remote zones observed with control CDC EV therapy (n=3 mice per group). Scale bar = 50 μm. *p<0.05 using a one-way ANOVA followed by Tukey’s post-hoc test. Data are represented as mean ± SEM.

### CDC EV miR cargo has a differential effect on infarct and remote zone apoptosis one-month post-MI

To examine the effect of EV therapy on apoptosis in the remote and infarct zone following ischemic injury, we analyzed cleaved caspase-3-stained sections from CDC EV, DROSHA EV and PBS treated hearts at 4 weeks post-MI. The percentage of apoptotic cells remained highest in the infarct region and lowest in the remote zone for all treatment groups at one month (**Fig 6**). Interestingly, both CDC EVs and DROSHA EVs demonstrated a significant reduction in the percentage of apoptotic cells in the infarct zone as compared with PBS control (CDC-EV: 10 ± 0.3%, DROSHA-EV: 12 ± 0.6%, PBS: 17 ± 0.6%). This suggests a role for additional non-miR cargo in mediating EV reduction of infarct zone apoptosis at 1-month post-MI (**Fig 6A-C,G**). In the remote zone, administration of CDC EVs resulted in a decrease in the % of cleaved caspase-3 cells when compared with PBS control (CDC EV: 1 ± 0.1%, DROSHA EV: 3 ± 0.9%, PBS: 2 ± 0.03%) (**Fig 6D-F,H**). (n=3 mice/group, mean ± SEM, *p<0.05 using one-way ANOVA with Tukey’s post-hoc test). In contrast, DROSHA EV treatment did not alter remote zone apoptosis and suggests that miR cargo was necessary for EV miR-mediated reduction of remote zone apoptosis. Together, these results demonstrate that miR cargo are vital for mediating several aspects of the therapeutic benefits provided by CDC-EV treatment.

**Figure 6.**
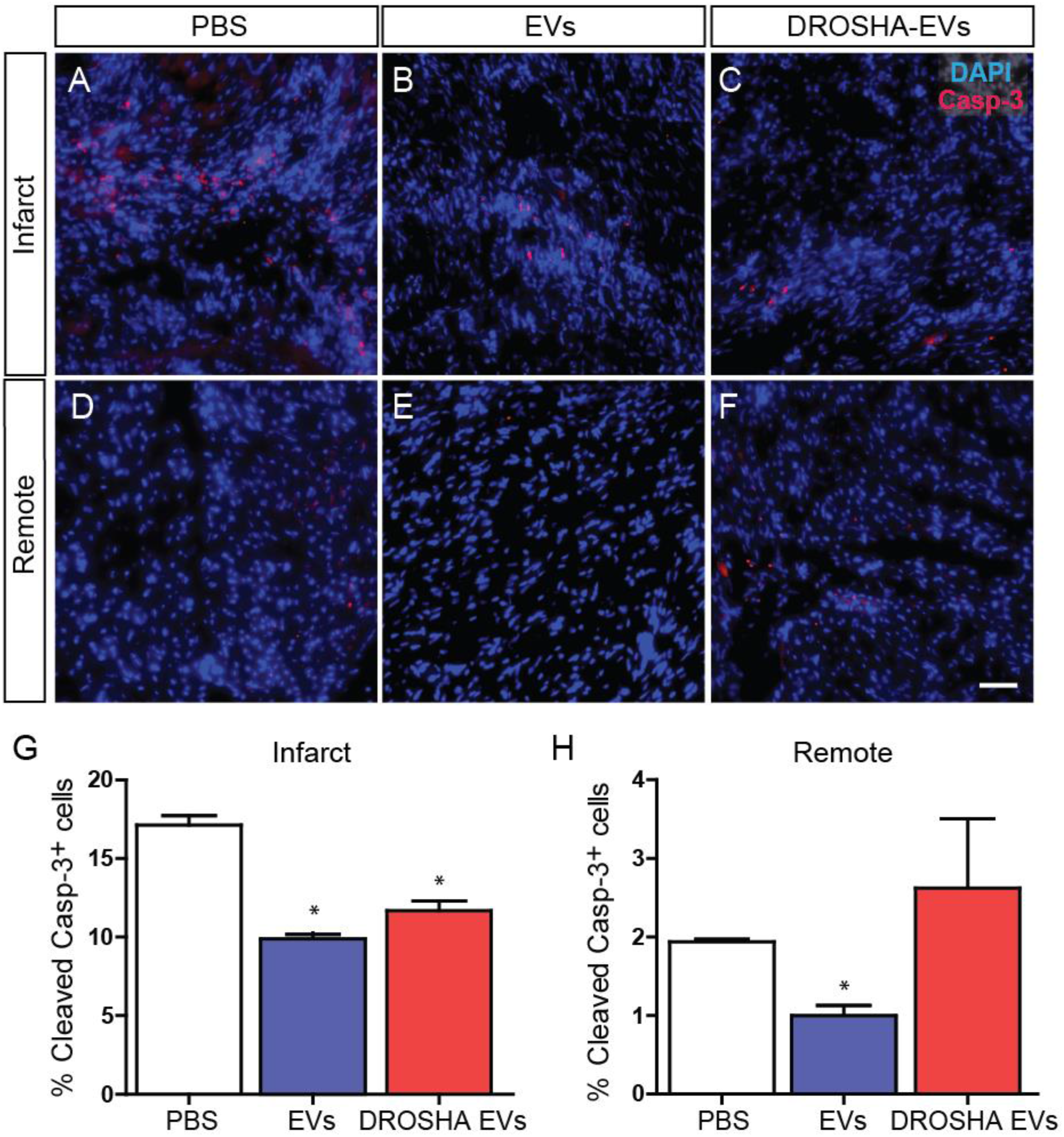
hCDC EV miR cargo differentially regulates cell death in the infarct vs. remote myocardium. Cleaved caspase-3-stained sections from the **(A-C)** infarct and **(D-F)** remote zone 28 days post-MI. **(G)** While control CDC EVs and DROSHA KD EVs both reduced the percentage of apoptotic cells in the infarct zone, **(H)** treatment with miR depleted DROSHA KD EVs failed to match CDC EVs in reducing apoptosis in the remote myocardium at the same time point. (n=3 hearts per group). Scale bar = 50 μm. *p<0.05 using a one-way ANOVA followed by Tukey’s post-hoc test. Data are represented as mean ± SEM.

### *In vitro* screening identifies synergistic candidate miRNAs as mediators of myocyte apoptosis and fibroblast proliferation

Having established the necessity of miR cargo for mediating the cellular effects of CVC-EV treatment *in vivo*, we hypothesized that specific combinations of miR cargo would be sufficient to impart therapeutic benefit on otherwise inert EVs. Using normal human dermal fibroblast (NHDF) that generate inert EVs, we performed direct transfection of three candidate miRs that were differentially expressed in CDCs. The contribution of miR-146a-5p, miR-126-3p and miR-370-3p were tested alone and in combination to determine their impact on cardiomyocyte apoptosis and fibroblast proliferation. Both primary neonatal mouse cardiomyocytes and NHDFs were transfected with media control, miR-146a mimics, miR-126 mimics, miR-370 mimics, or the combination of 2 or 3 at 1/2 or 1/3 dose, respectively. Assessing the percentage of TUNEL^+^ myocytes following miR mimic transfection, we saw significant main effects of both miR-146a (F(1,24)=26.2, p<0.001) and miRNA 370 (F(1,24)=31.2, p<0.001), indicating that both miRNAs played an important role in cardiomyocyte apoptosis. Furthermore, specific comparisons between groups revealed larger and more significant effects of miRs in combination (Dunnett’s multiple comparisons test) (**Fig 7A-C**). We found that miR-370 synergizes separately with miR-126 (p=0.0058) and miR-146a (p=0.0003) to yield enhanced anti-apoptotic effects on cardiomyocytes when compared with each miR in isolation and, importantly, the combination of all three miRNAs resulted in the greatest reduction in myocyte apoptosis (p<0001) (n=4, mean ± SEM, *p<0.05 using three-way ANOVA with Dunnett’s multiple comparisons test).

**Figure 7.**
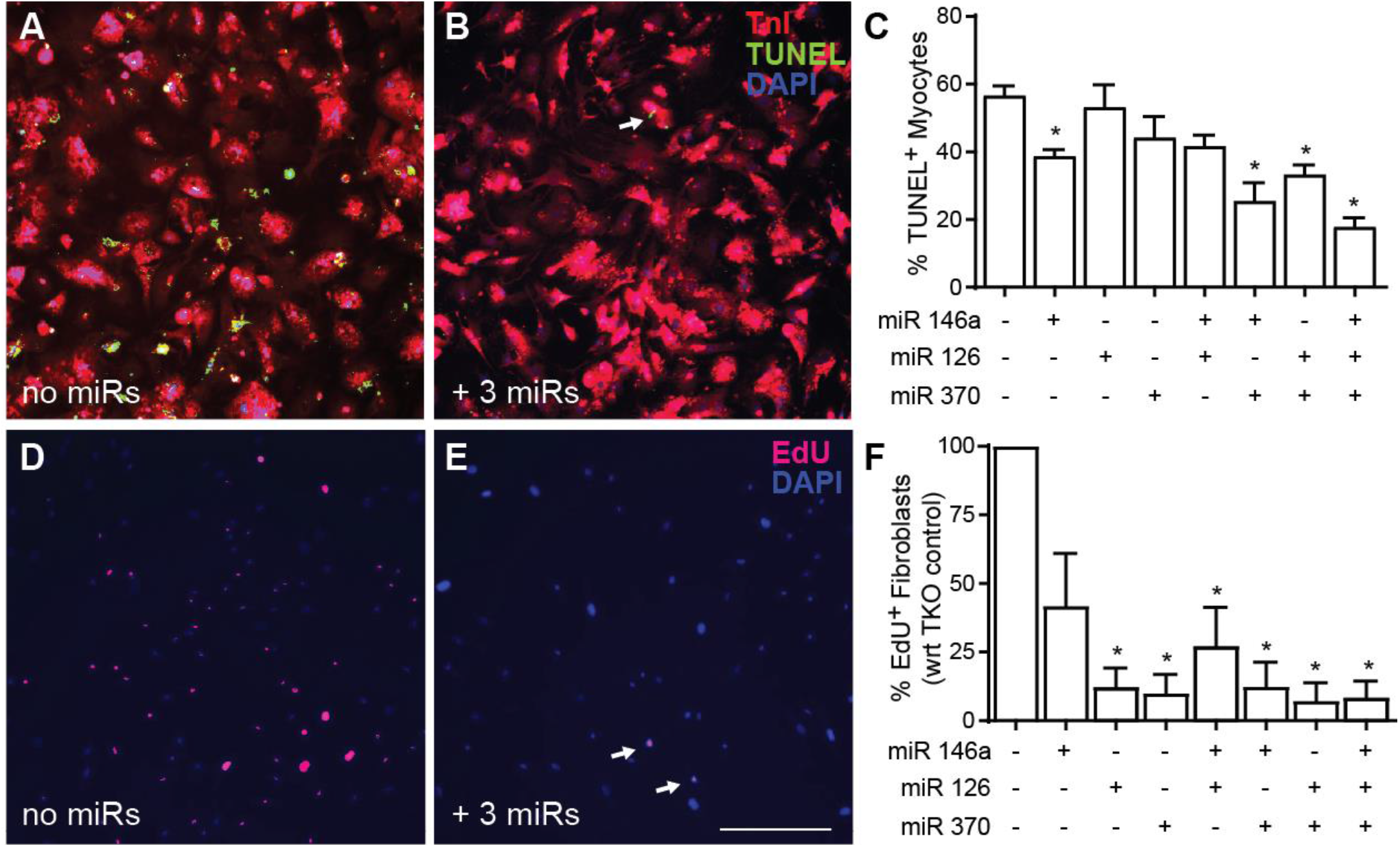
*In vitro* screening by direct target cell transfection identifies synergistic candidate miRNAs mediating myocyte apoptosis and fibroblast proliferation. **(A-C)** Direct transfection of primary neonatal mouse cardiomyocytes with a combination of all three candidate miRs was the most efficacious paradigm for reducing TUNEL-positive nuclei at one week in an *in vitro* assay of apoptosis (n=4). **(D-F)** Quantification of the percentage of EdU^+^ fibroblasts (5-hour pulse) following direct transfection of candidate miR combinations identified miR-126 and miR-370 as highly significant modifiers of fibroblast proliferation *in vitro* (n=3). Scale bar = 400 μm. *p<0.05 using a three-way ANOVA followed by Dunnett’s post-hoc test. All data are represented as mean ± SEM.

Assessing the percentage of EdU^+^ fibroblasts following miR mimic transfection, we saw significant main effects of miR-126 and miR-370 (n=3, *p<0.05 using three-way ANOVA with Dunnett’s multiple comparisons test) (**Fig 7D-F**). There was no significant effect of miR-146a alone on fibroblast proliferation (n=3, p=0.1 using three-way ANOVA with Dunnett’s multiple comparisons test), nor was there any significant synergistic effect between miRs.

### The combination of miR-146a, 126, and 370 were sufficient for EV-mediated reduction of cardiomyocyte apoptosis *in vitro*

As reduction of cardiomyocyte apoptosis has been linked with improvement in systolic function and scar size post-MI, we sought to determine if the combination of miRs that resulted in the lowest level of cardiomyocyte apoptosis following direct mimic transfection were sufficient to rescue myocyte cell death when delivered in combination by EVs. We performed a gain-of-function study on therapeutically inert NHDFs, transfecting them with a 1:1:1 ratio of miR-146a, miR-126 and miR-370 mimics. Isolated NHDF EVs demonstrated comparable levels of miR-146a-5p, miR-126-3p, and miR-370-3p as CDC EVs. While unmodified NHDF EVs had no significant effect on preventing myocyte cell death from apoptosis compared with PBS control at one week, NHDF EVs enriched in therapeutic miRs demonstrated a significant reduction in cardiomyocyte cell death that was comparable with that of CDC EVs, highlighting the sufficiency of this combination of EV delivered miRs to rescue cardiomyocyte apoptosis. (n=7, mean ± SEM, *p<0.05 using one-way ANOVA with Tukey post-hoc test) (**Fig 8**).

**Figure 8.**
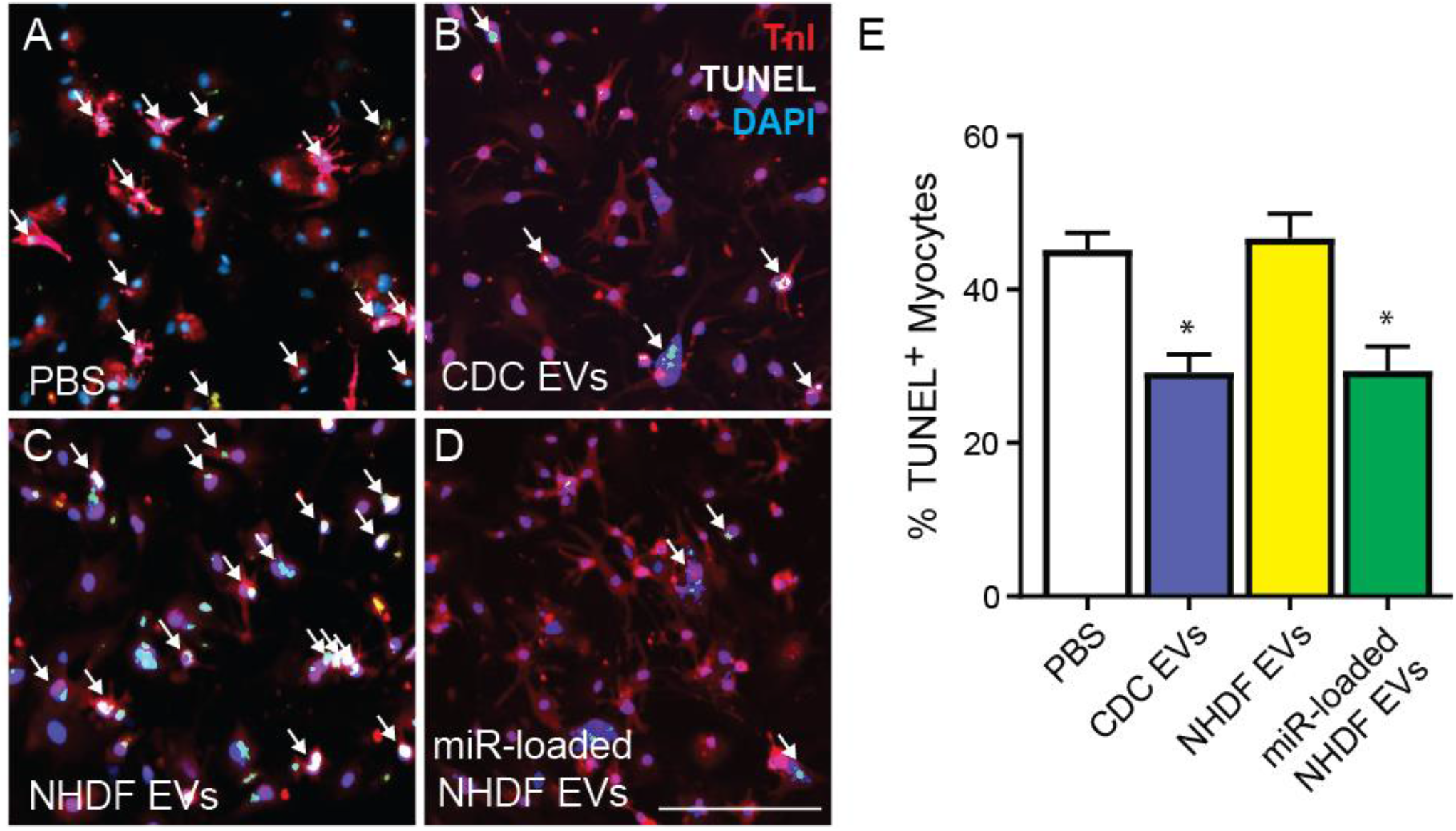
The combination of miR 146a, 126, and 370 are sufficient for EV-mediated reduction of cardiomyocyte apoptosis *in vitro*. Primary neonatal mouse cardiomyocytes were treated for 24 hours with **(A)** PBS control, **(B)** CDC EVs, **(C)** normal human dermal fibroblast (NHDF) EVs, or **(D)** NHDF EVs loaded with miRs-146a, −126, and −370 and stained for TnI, DAPI and TUNEL to quantify % myocyte apoptosis. **(E)** The combination of miR-146a, miR-126 and miR-370 are sufficient to enable inert NHDF EVs to rescue cardiomyocyte apoptosis at the level of CDC-EVs. Scale bar = 100μm. *p<0.05 using a two-way ANOVA followed by Tukey’s post-hoc test. All data are represented as mean ± SEM.

## Discussion

Extracellular vesicles are important intercellular delivery vehicles released by cells which facilitate the transfer of proteins, small RNAs, and lipids, with roles in both steady-state physiology and disease pathology. EVs isolated from specific cell populations, including CDCs, have demonstrated therapeutic efficacy in pre-clinical models of ischemic heart disease, highlighting them as promising vectors for the treatment of cardiovascular disease (4–10). They have been shown to regulate fibrosis (20), angiogenesis (21), cardiomyocyte hypertrophy (22), apoptosis (23), and the immune response (24) following injury. In addition, CDC-derived EVs have also demonstrated the ability to impart their therapeutic properties to inert cells, rendering skin fibroblasts therapeutic when preconditioned by CDC-EVs in a rat model of MI (25). Given the heterogeneity of their cargo and diversity of their responses, a greater understanding of the specific biological components necessary for their intrinsic activity is needed to realize the potential of EV-based therapeutics.

In this study, we demonstrate that EV-delivered miRNAs play a critical role in the cardioprotective properties of human CDC-EVs in an *in vivo* model of AMI. We observed that knockdown of Drosha in CDCs globally reduced miRNA cargo in CDC-EVs without changing expression of EV surface markers or protein cargo. This was associated with a reduction of *in vitro* metrics of efficacy, including angiogenesis, cardiomyocyte apoptosis and fibroblast proliferation, as well as a significant reduction in cardiac function with respect to wild-type hCDC-EVs. The crucial role of EV miRNAs following ischemic cardiac injury was observed not only at a functional level but also at a cellular level *in vivo*. In contrast to wild-type hCDC-EVs, Drosha-EVs failed to regulate remote interstitial fibrosis, macrophage density, and remote zone apoptosis following AMI, highlighting the diversity of biological responses mediated by hCDC-EV shuttled miRNAs.

miRNAs are a class of small non-coding RNAs that function as endogenous mediators of RNA silencing. While they are the most frequently studied extracellular RNA, they encompass only a minor element of RNA biotypes in EVs. Additional RNA biotypes, such as long non-coding RNA (lncRNA), small nucleolar RNA (snoRNA), small nuclear RNA (snRNA) and Y RNA, likely constitute a larger fraction in EV cargo (26, 27). Elevated content of Y RNA fragment EV-YFI within CDC-EVs, for example, has been shown to correlate with *in vivo* CDC potency (28). Nevertheless, prior studies have demonstrated cardioprotective effects of individual miRs in models of disease, suggesting their functional importance. Treatment of neonatal rat cardiomyocytes with a miR-146a mimic increased viability and protected cardiomyocytes from oxidative stress, potentially mediated by downregulation of IRAK1 and TRAF6 (29). Ischemia and oxidative stress induced by hydrogen peroxide treatment of H9C2 cells was reversed following treatment with a miR-370 mimic, along with a reduction in cardiomyocyte apoptosis, potentially mediated by the downregulation of transcription factor FOXO1 (30). miR-24 attenuated fibrosis in the infarct border zone two weeks post-MI (31) and miRs 125a and 126 have been shown to promote angiogenesis (32, 33). *In vivo* studies have also demonstrated immunomodulatory capabilities of EV miRNA cargo, with CDC-EV delivered miR-181b implicated in reducing macrophage infiltration into the myocardium following I/R and decreasing expression of NOS2 and TNF proinflammatory genes in cardiac macrophages (34). Despite the implicated role of individual miRNAs in cardioprotective responses, until now no prior study has established that endogenous miRNA are responsible and necessary for mediating the therapeutic benefits of human CDC-EVs following AMI.

Surprisingly, treatment with mouse CDC-EVs *in vitro* failed to recapitulate the cardioprotective effects of human CDC-EVs in assays of angiogenesis, proliferation and cell death, despite their surface marker similarity. These results appear to be related to differences in bioactive miRNA cargo between mouse and human CDC-EVs, with evidence of several highly expressed hCDC-EV miRs absent in their murine counterparts. Differences in EV miRNA content between species may be secondary to differences in the physiological state of the parent cell (human CDCs are obtained from tissue of patients undergoing on-pump CABG, while mouse CDCs are harvested from healthy mice) or differences in the mechanisms that control nucleic acid sorting into human and mouse EVs (35). This outcome highlights the potential pitfalls of extrapolating studies conducted with animal EVs to human responses. Additional work is necessary to further elucidate the individual contributions of species-specific effects vs. parent cell physiological state in influencing miRNA cargo between mouse and human CDCs-EVs.

We hypothesized that a cluster of differentially regulated miRs was responsible for mediating the overall cardioprotective effects of hCDC-EV miRNA cargo. We identified three candidate miRs, differentially expressed in CDC-EVs vs. inert NHDF-EVs, that have been shown to inhibit the RNA expression levels of genes regulating apoptosis and autophagy (36–38). We demonstrated that both direct transfection and EV loading of the candidate miRs resulted in differential effects on cardiomyocyte apoptosis and fibroblast proliferation. Perhaps most striking was the synergistic regulatory effects of miR-146a, miR-370 and miR-126 on the reduction of myocyte apoptosis following direct transfection. Furthermore, when loaded into therapeutically inert dermal fibroblast EVs, this miR cluster rendered NHDF-EVs capable of recapitulating the cardioprotective effects of CDC-EVs on cardiomyocyte viability. Prior studies investigating the therapeutic role of EV-associated miRNA clusters provide greater insight on this phenomenon, whereby groups of miRNAs have been shown to work in concert to elicit a therapeutic benefit. For example, by simultaneously targeting different genes of the Notch3 signaling pathway, the EV-associated miR-106a-363 cluster enhanced LV ejection fraction in a mouse model of ischemic heart injury (39). In contrast to miRNA clusters which regulate genes along the same molecular pathway, miR-146a-370-126 appears to target independent apoptosis- and fibrosis-associated pathways (30, 37, 40, 41). This may account for their combined effect on reducing cardiomyocyte viability when administered in concert. Future network-based studies focused on miRNA synergism may provide useful information on miRNA functions at a systems level (42).

While this study focused on understanding the *endogenous* cargo responsible for CDC-EV mediated cardioprotection, it also demonstrated the applicability of *exogenous* miR cargo loading in transforming inert EVs into therapeutic vectors. Additional studies in other disease models have investigated the controlled loading of miRNAs into EVs and highlight the potential for clinical translation of this engineering strategy. For example, EVs loaded with antitumor miRNAs (miR-31 and miR-451a) significantly enhanced hepatocellular carcinoma apoptosis as compared with control EVs (43). Exogenous loading of miRNAs directly into EVs has also been explored as an alternate strategy for improved EV therapeutic efficacy. Didiot *et al* demonstrated that EVs produced by glioblastoma U87 cells could be directly loaded with hydrophobically modified siRNA which resulted in enhanced silencing of target genes (44). Various additional techniques have been explored to load purified EVs with therapeutic miRNA cargo, including electroporation, sonication, and saponin, but require further studies to ensure the proper loading of miRNA *into* EVs and at the same time elucidate miRNA loading efficiency (45).

In conclusion, the results of this study demonstrate that miRNAs play a critical role in the regenerative potential of CDC-derived extracellular vesicles. The global downregulation of EV miRNA cargo attained by Drosha knockdown in human CDCs abrogated the ability of hCDC-EVs to promote recovery following acute MI. Furthermore, the combination of miR-146a-5p, miR-370-3p and miR-126a-3p demonstrated synergy and sufficiency in rescuing cardiomyocyte apoptosis. These results identify a potential therapeutic miR cluster for rescuing ischemia induced apoptosis and highlight the potential for tailoring EV miR cargo to further enhance cardiac regeneration after myocardial infarction.

## Acknowledgements

We would like to thank Dr. Min Gao from Kent State for technical assistance with cryo-TEM, Beth Palka for technical assistance in murine Drosha primer design and Rebeccah Young for technical assistance with mouse CDC characterization.

## Abbreviations

AMI: acute myocardial infarction
CDC: cardiosphere-derived cell
CM: cardiomyocyte
CMP: cardiomyocyte specific peptide
EF: ejection fraction
EVs: extracellular vesicles
HUVEC: human umbilical vein endothelial cell
lncRNA: long non-coding RNA
LV: left ventricle
MI: myocardial infarction
miR: microRNA
NHDF: normal human dermal fibroblast
NTA: Nanoparticle Tracking Analysis
snRNA: small nuclear RNA
snoRNA: small nucleolar RNA

---

**Supplementary Figure 1.**
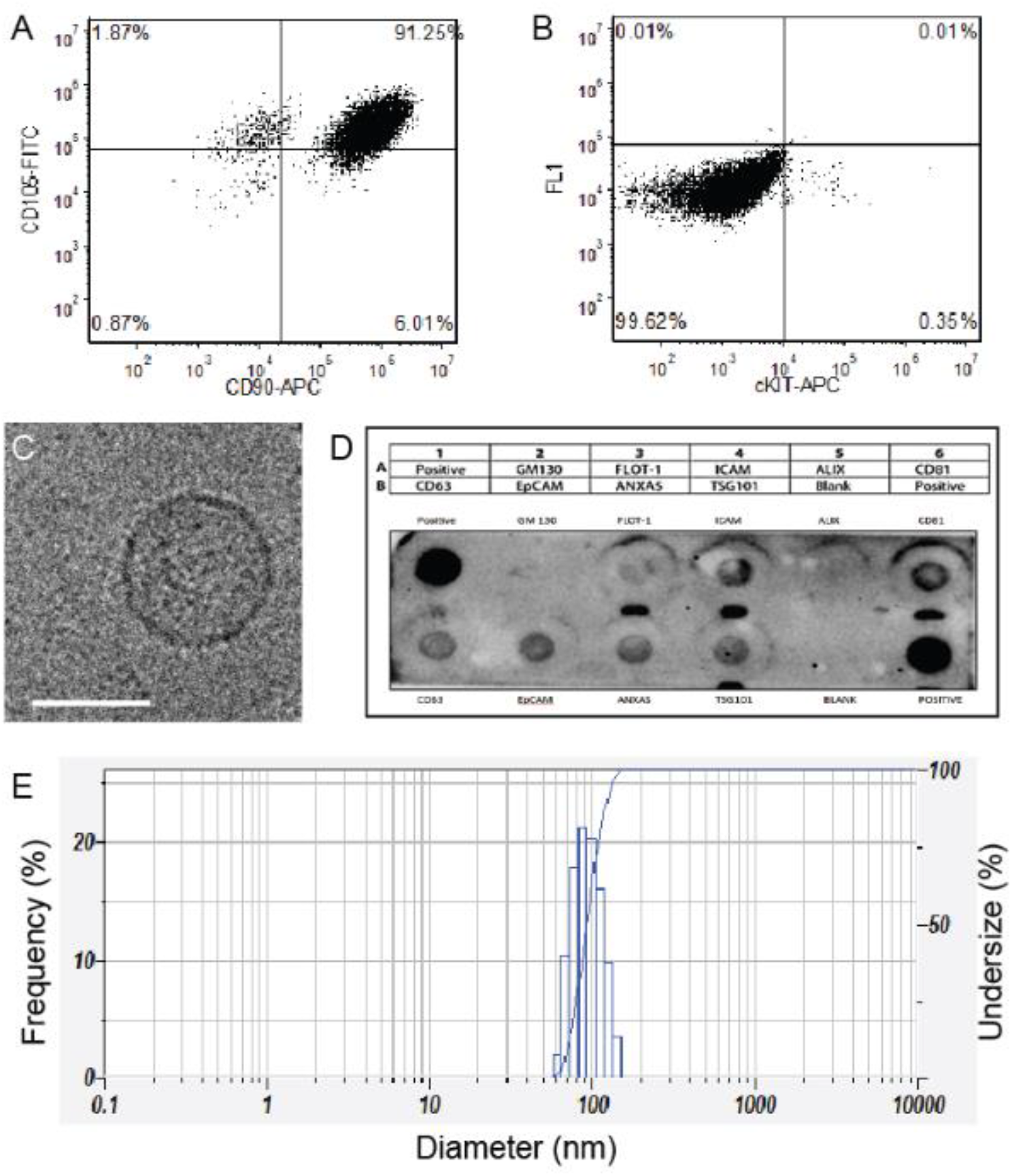
Characterization of mouse cardiosphere-derived cells and mCDC EVs. **(A-B)** Representative flow cytometry plots of mCDCs stain with CD90, CD105 and cKIT. mCDCs express high levels of mesenchymal markers CD90 and CD105 with low levels of cKIT, like their human counterparts. **(C)** Cryo-TEM image of mCDC EVs isolated from serum free tissue culture. Scale bar: 100μM. **(D)** Exo-Check Exosome Antibody Array for exosomal surface protein marker detection and assessment of cellular contamination of mCDC EVs.

**Supplementary Figure 2:**
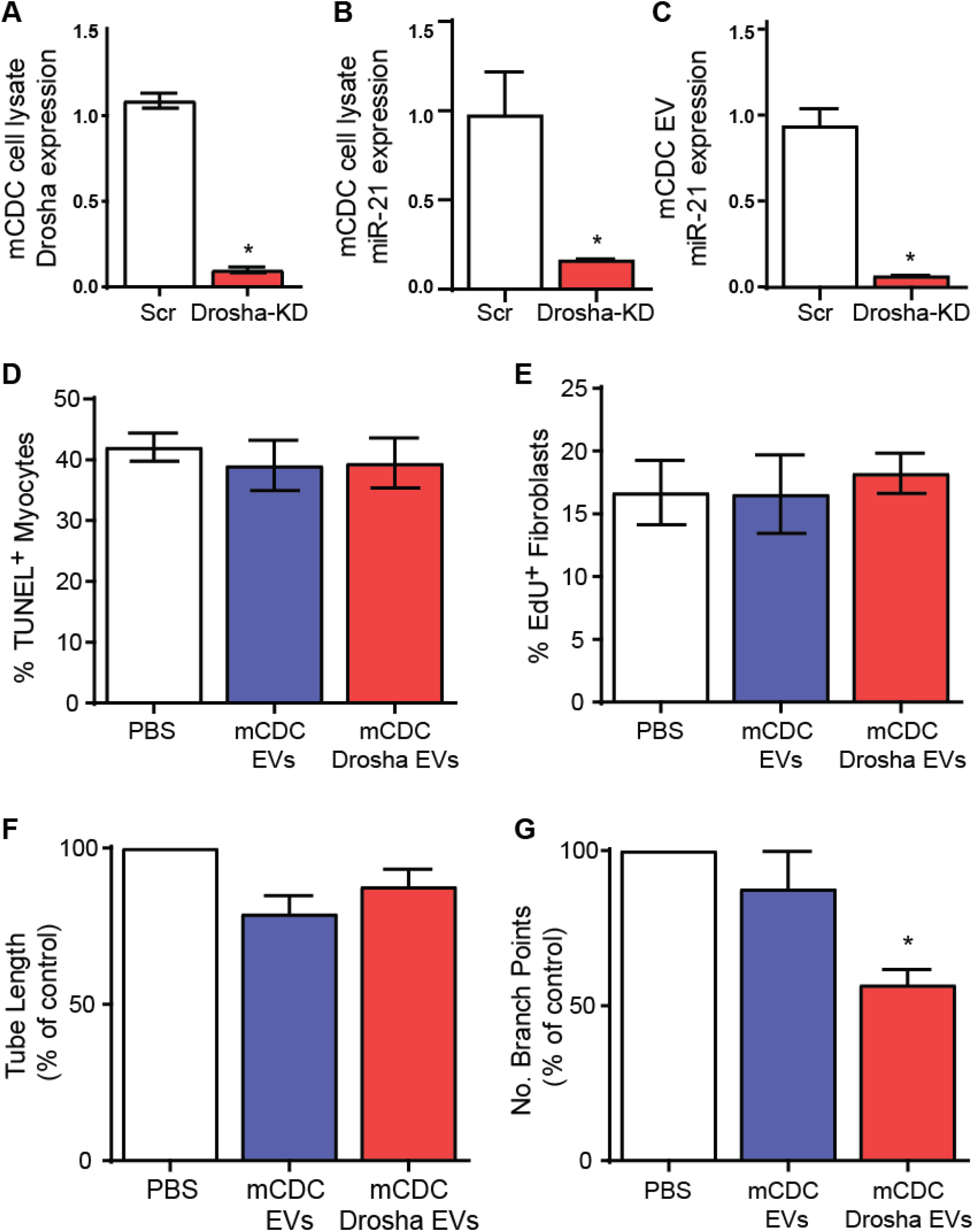
Characterization of DROSHA siRNA transfected mouse CDCs and their miR depleted EVs. **(A)** Relative expression of DROSHA mRNA in mCDCs transfected with Scr or Drosha siRNA. Relative expression of candidate miRNA (miR-21) in **(B)** mCDCs transfected with Scr and DROSHA siRNA and **(C)** mCDC-EVs isolated from Scr and DROSHA siRNA transfected cells (n=3). *p<0.0001 using unpaired student’s t-test. Mouse CDC EVs were unable to recapitulate human CDC EVs in their ability to **(D)** reduce terminal deoxynucleotidyl transferase nick end labeling (TUNEL)-positive cardiomyocytes (CMs) in an *in vitro* apoptosis assay (n=8), **(E)** reduce fibroblast proliferation as assessed by EdU staining with a 5-hour pulse (n=4), and **(F-G)** stimulate endothelial migration and angiogenesis *in vitro* (n=7), demonstrating matched levels with PBS control (n=7). p>0.05 using one-way ANOVA with Tukey’s post-hoc test. miR depleted mCDC EVs showed no appreciable change with respect to mCDC EVs or PBS control with exception to number of branch points in an endothelial tube assay in vitro (n=7) *p<0.05 using one-way ANOVA with Tukey’s post-hoc test. All data represented as mean ± SEM.

**Supplementary Figure 3:**
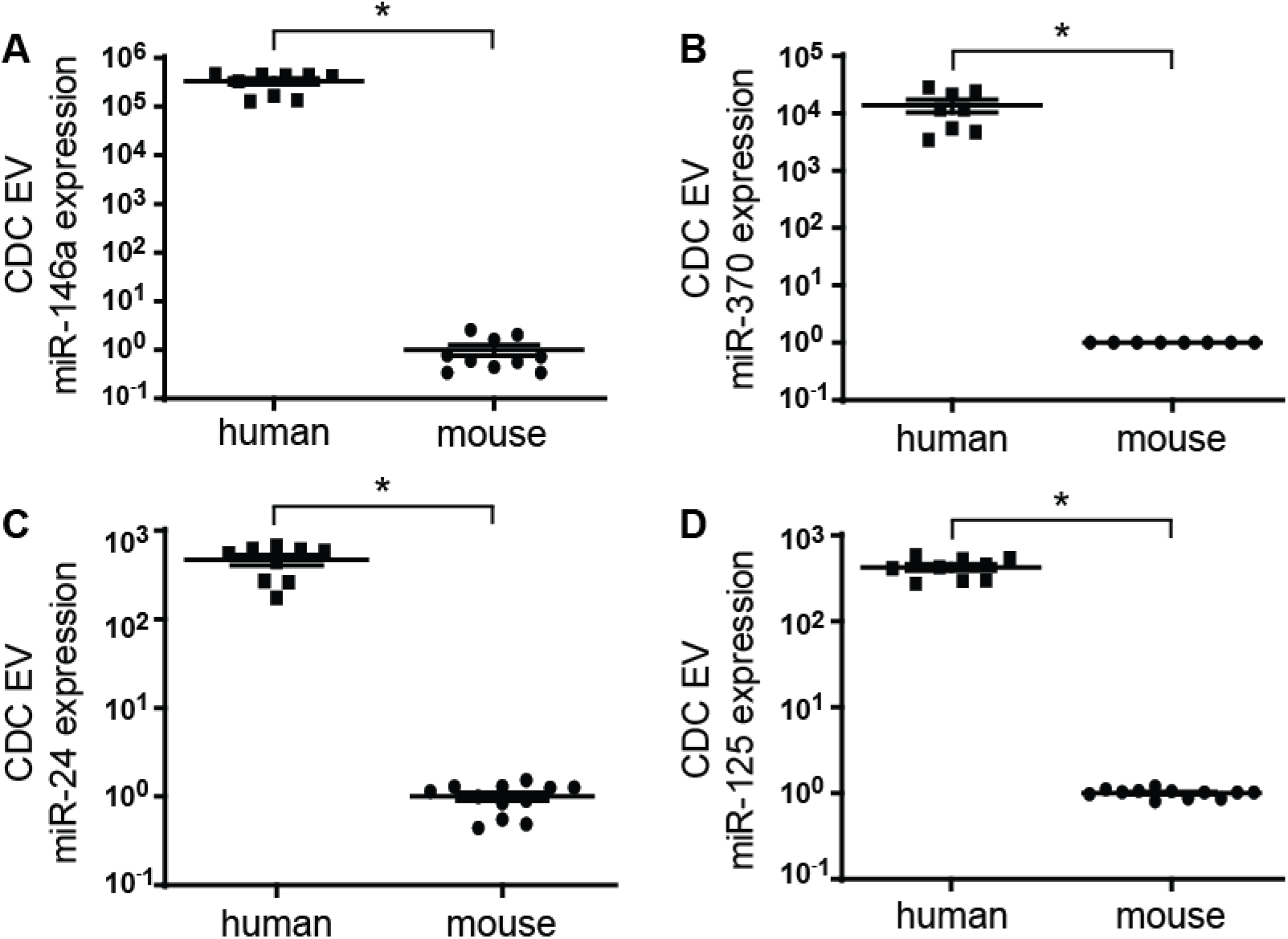
mCDC EVs lack potentially therapeutic miRNAs seen to be upregulated in hCDC EVs. Relative expression of **(A)** miR-146a, **(B)** miR-24, **(C)** miR-125, and **(D)** miR-370 in mouse and human CDC EVs with respect to mCDC EVs (n=8-12). *p<0.0001 using an unpaired student’s t test. All data are represented as mean ± SEM.

